# A deep generative model for capturing cell to phenotype relationships

**DOI:** 10.1101/2024.08.07.606396

**Authors:** Ethan Weinberger, Patrick Yu, Su-In Lee

## Abstract

Single-cell omics has proven to be a powerful instrument for exploring cellular diversity. With advances in sequencing protocols, single-cell studies are now routinely collected from large-scale donor cohorts consisting of samples from hundreds of donors with the goal of uncovering the molecular bases of higher-level donor phenotypes of interest. For example, to better understand the mechanisms behind Alzheimer’s disease, recent studies with up to hundreds of samples have investigated the relationships between single-cell omics measurements and donors’ neuropathological phenotypes (e.g. Braak staging) [4, 9, 3, 10].

In order to ensure the robustness of such findings, it may be desirable to aggregate data from multiple distinct donor cohorts. Unfortunately, doing so is not always straightforward, as different cohorts may be equipped with different sets of phenotype labels. Continuing the previous Alzheimer’s example, recent AD study cohorts have provided various subsets of neuropathological phenotypes, cognitive testing results, and *APOE* genotype. Thus, it is desirable to be able to infer any missing phenotype labels such that all available cell-level data in the study of a given phenotype of interest could be used. Moreover, beyond simply imputing missing phenotype information, it is often of interest to understand which groups of cells and/or molecular features may be most predictive of a given phenotype of interest. As such, there is a pressing need for computational methods that can connect cell-level measurements with donor-level labels.

However, accomplishing this task is not straightforward. While a rich literature exists on learning meaningful low-dimensional representations of cells [7, 8, 1, 2] and for inferring corresponding cell-level labels (e.g. cell type) [11], the donor level prediction task introduces substantial additional complexity. For example, different numbers of cells may be recovered from each donor, and thus our prediction model must be able to handle arbitrary numbers of samples as input. Moreover, ideally our model would not *a priori* require any additional prior knowledge beyond our cell-level measurements, such as the importance of different cell types for a given prediction task.

To resolve these issues, here we propose milVI (multiple instance learning variational inference), a deep generative modeling framework that explicitly accounts for donor-level phenotypes and enables inference of missing phenotype labels post-training. In order to handle varying numbers of cells per donor when inferring phenotype labels, milVI leverages recent advances in multiple instance learning. We validated milVI by applying to impute held-out Braak staging information from an Alzheimer’s disease study cohort from Mathys et al. [9], and we found that our method achieved lower error on this task compared to naive imputation methods.

## 1 The milVI model

For a given donor *b*, we first draw a phenotype label

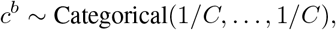

where *C* denotes the number of possible phenotype labels. Within a bag, for a cell *i* we first draw a low-dimensional representation

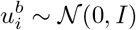

representing phenotype-agnostic variations. We then transform 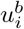 along with donor phenotypes *c*^*b*^ to the parameters of a second set of latent variables

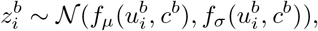

representing phenotype-aware variations. Similar to previous works in generative modeling of scRNA-seq data [7, 11, 2], we then transform 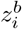 into the parameters of a negative binomial distribution to generate our observed data. Our generative model bears resemblance to that of the scANVI method of Xu et al. [11]. However, rather than assuming the presense of cell-level labels, milVI instead assumes that labels represent donor-level information. We provide a graphical representation of our model in Figure 1b.

**Figure 1:**
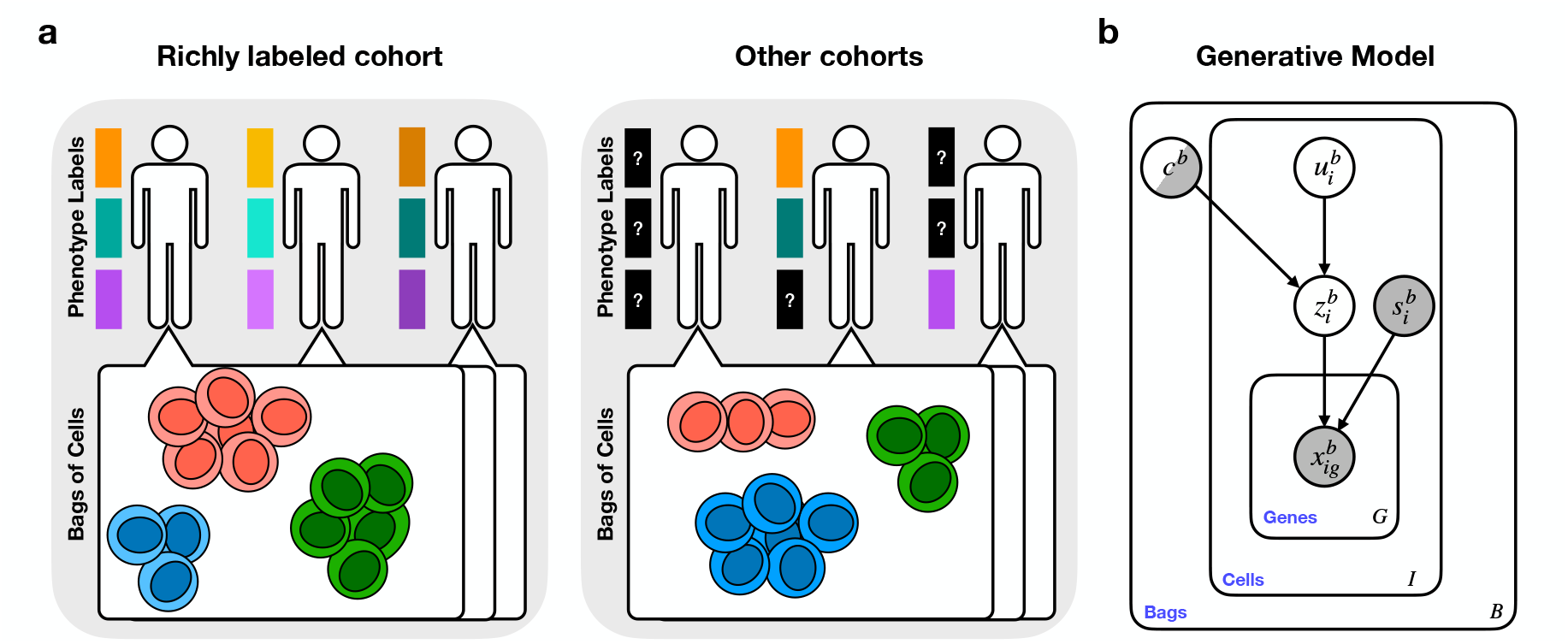
**a**, Modern single-cell analyses often consider data from multiple distinct cohorts of donors. While some cohorts may be equipped with rich phenotype information, others may be missing phenotype labels for some or all patients. **b**, The milVI model. Shaded nodes denote observed variables, while half-shaded nodes indicate variables that are only sometimes observed. Arrows denote conditional dependence, and plates indicate independent replication.

With our model in hand, we now specify a procedure for inference. As the posterior distribution of our model is intractable, we use variational inference to perform approximate inference over our latent variables. For a donor *b*, we assume that our variational distribution factorizes as:

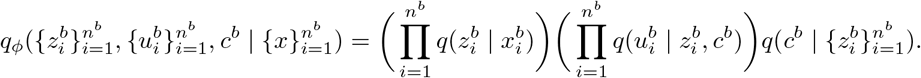

Following Kingma et al. [6] we may then derive two evidence lower bounds (ELBOs): one in the case of an observed phenotype label *c*^*b*^, and the other in the case of an unobserved label. For the most part (Appendix A), these derivations use standard machinery established in [6] and [11]. However, we draw special attention to the term 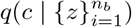 in our variational distribution, which must be treated with care due to the conditioning on arbitrary numbers of cell embeddings. To accommodate this constraint, we leverage techniques from the multiple instance learning literature. In particular, we make use of the attention-based multiple instance learning (ABMIL) framework of Ilse et al. [5].

The ABMIL mechanism pools a bag of embeddings via a weighted average embedding ∑_*i*_ *a*_*i*_*z*_*i*_, the result of which can then be fed to standard feedforward neural networks. To enable the model to dynamically infer the importance of each instance for a given prediction, the weights of this average are determined by an attention mechanism

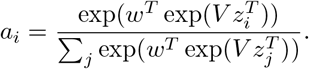

To implement 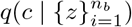, we thus first pool the given *z*’s for a bag via the attention mechanism, and then pass the resulting weighted average embedding into a feedforward neural network to infer the parameters of the categorical distribution.

## 2 Results

To validate milVI’s performance, we measured its ability to predict held-out Braak neuropathology scores from scRNA-seq data from the prefrontal cortex collected by Mathys et al. [9]. In particular, we randomly held out Braak labels from 20% of patients during training and predicted their Braak labels during evaluation. To assess the quality of milVI’s predictions, we calculated the mean absolute error (MAE) as well as Spearman and Pearson correlations between the held-out patients’ true and predicted Braak labels. We benchmarked milVI against two naive prediction pipelines. First, we trained logistic regression models to predict Braak scores using pseudobulk expression profiles from each donor. Second, we trained logistic regression models to predict Braak scores using cell-level data; cell-level predictions were then averaged to obtain final donor-level predictions. We found that milVI outperformed both baselines (Figure 2). However, all methods exhibited relatively high variance in performance across train-test splits, potentially due to overfitting given the small number (48) of total donors. For future work we will investigate milVI’s generalization performance after training on larger-scale donor cohorts, as well as investigate milVI’s capabilities to facilitate new biological insights (e.g. via inspection of the cell-level attention weights).

**Figure 2:**
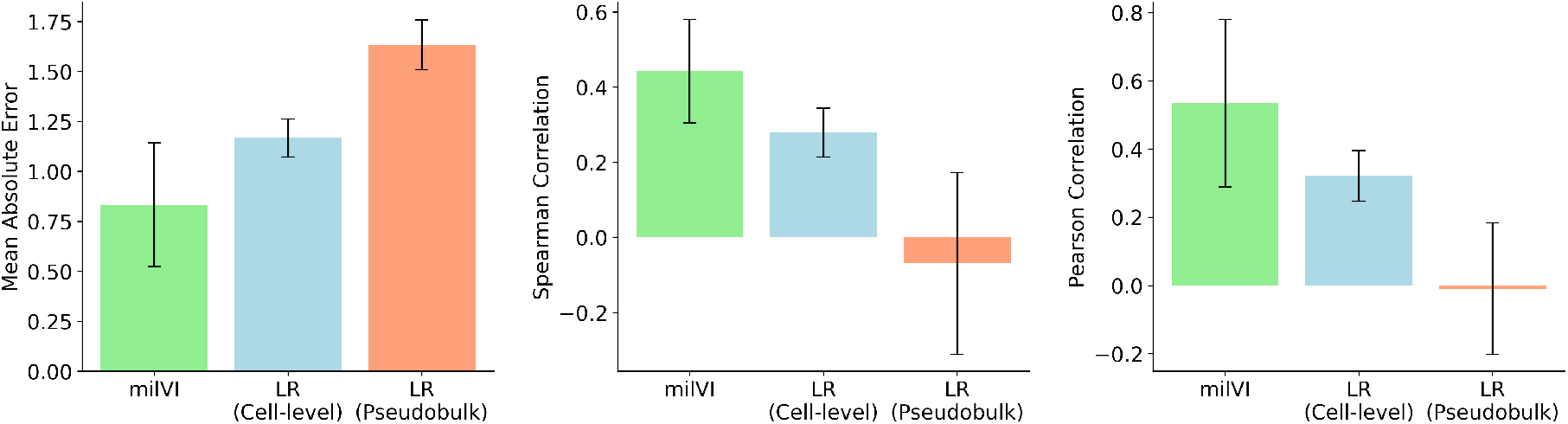
Assessing milVI and baseline models’ performance at predicting held-out Braak neuropathology scores. Plots depict mean ± standard deviation across three train-test splits for each metric.

## A Derivation of milVI evidence lower bounds

Here we provide derivations of the evidence lower bound (ELBO) used to train milVI. The below derivation largely follows that of Xu et al. [11], with corresponding changes to accommodate bag-level labels as opposed to cell-level labels. We begin by considering the unlabelled cell case. As noted in the main text, for a given bag *b* we assume that our variational distribution factorizes as

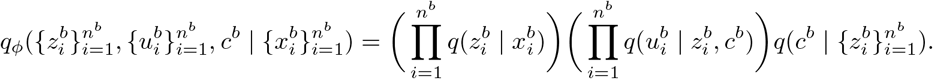

Using Jensen’s inequality and weighting by our variational distribution 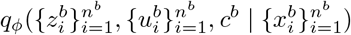 we have:

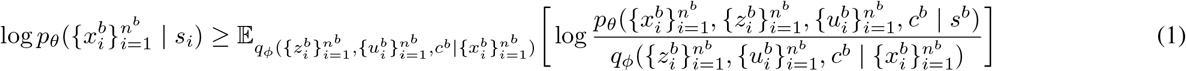

Assuming that all cells within a bag are generated i.i.d. we can rewrite the right-hand side of Equation 1 as

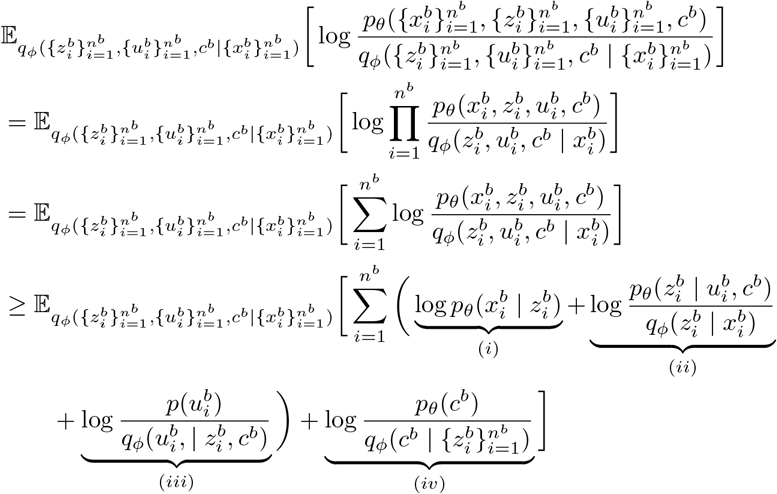

We now proceed to simplify the terms in the preceding inequality as follows. For term (*i*) we have:

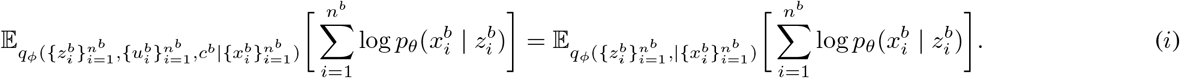

For term (*ii*) we have:

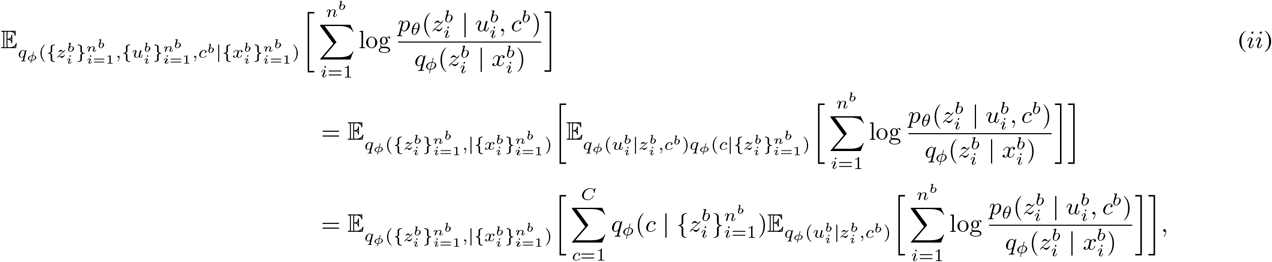

where here the sum over *C* indexes over possible bag labels.

For term (*iii*) we have:

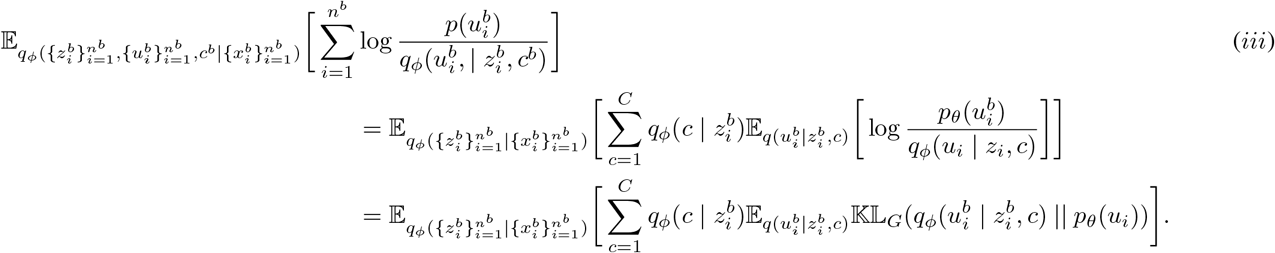

For term (*iv*) we have:

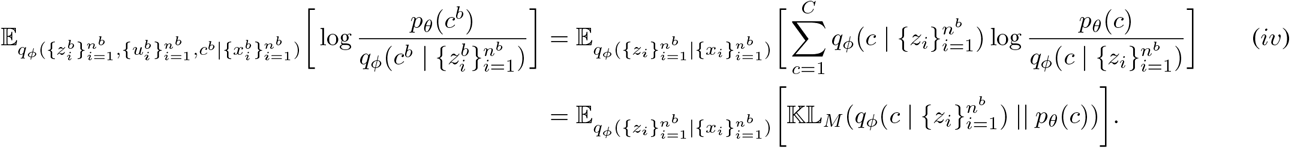

where 𝕂𝕃_*M*_ and 𝕂𝕃_*G*_ denote multinomial and Gaussian KL divergences.

Now we consider the case where bag label information is available. In this case the resulting bound is nearly identical to that of the unsupervised case, with the addition of an extra classification loss term with weight *α* that encourages the posterior 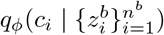 to match the provided label information.

## Notes

https://www.nature.com/articles/s41586-019-1195-2

### Competing Interest Statement

The authors have declared no competing interest.

## References

[1] Matthew Amodio, David Van Dijk, Krishnan Srinivasan, William S Chen, Hussein Mohsen, Kevin R Moon, Allison Campbell, Yujiao Zhao, Xiaomei Wang, Manjunatha Venkataswamy, et al. Exploring single-cell data with deep multitasking neural networks. Nature methods, 16(11):1139–1145, 2019.

[2] Gökcen Eraslan, Lukas M Simon, Maria Mircea, Nikola S Mueller, and Fabian J Theis. Single-cell rna-seq denoising using a deep count autoencoder. Nature communications, 10(1):390, 2019.

[3] Masashi Fujita, Zongmei Gao, Lu Zeng, Cristin McCabe, Charles C White, Bernard Ng, Gilad Sahar Green, Orit Rozenblatt-Rosen, Devan Phillips, Liat Amir-Zilberstein, et al. Cell subtype-specific effects of genetic variation in the alzheimer’s disease brain. Nature Genetics, 56(4):605–614, 2024.

[4] Mariano I Gabitto, Kyle J Travaglini, Victoria M Rachleff, Eitan S Kaplan, Brian Long, Jeanelle Ariza, Yi Ding, Joseph T Mahoney, Nick Dee, Jeff Goldy, et al. Integrated multimodal cell atlas of alzheimer’s disease. bioRxiv, pages 2023–05, 2023.

[5] Maximilian Ilse, Jakub Tomczak, and Max Welling. Attention-based deep multiple instance learning. In International conference on machine learning, pages 2127–2136. PMLR, 2018.

[6] Durk P Kingma, Shakir Mohamed, Danilo Jimenez Rezende, and Max Welling. Semi-supervised learning with deep generative models. Advances in neural information processing systems, 27, 2014.

[7] Romain Lopez, Jeffrey Regier, Michael B Cole, Michael I Jordan, and Nir Yosef. Deep generative modeling for single-cell transcriptomics. Nature methods, 15(12):1053–1058, 2018.

[8] Mohammad Lotfollahi, Mohsen Naghipourfar, Malte D Luecken, Matin Khajavi, Maren Büttner, Marco Wagenstetter, Žiga Avsec, Adam Gayoso, Nir Yosef, Marta Interlandi, et al. Mapping single-cell data to reference atlases by transfer learning. Nature biotechnology, 40(1):121–130, 2022.

[9] Hansruedi Mathys, Jose Davila-Velderrain, Zhuyu Peng, Fan Gao, Shahin Mohammadi, Jennie Z Young, Madhvi Menon, Liang He, Fatema Abdurrob, Xueqiao Jiang, et al. Single-cell transcriptomic analysis of alzheimer’s disease. Nature, 570(7761):332–337, 2019.

[10] Hansruedi Mathys, Zhuyu Peng, Carles A Boix, Matheus B Victor, Noelle Leary, Sudhagar Babu, Ghada Abdelhady, Xueqiao Jiang, Ayesha P Ng, Kimia Ghafari, et al. Single-cell atlas reveals correlates of high cognitive function, dementia, and resilience to alzheimer’s disease pathology. Cell, 186(20):4365–4385, 2023.

[11] Chenling Xu, Romain Lopez, Edouard Mehlman, Jeffrey Regier, Michael I Jordan, and Nir Yosef. Probabilistic harmonization and annotation of single-cell transcriptomics data with deep generative models. Molecular systems biology, 17(1):e9620, 2021.

